# Brown Bears activates a Satiety Hormone Cholecystokinin (CCK) pathway in adipose tissue during hibernation

**DOI:** 10.1101/2024.12.20.629672

**Authors:** T.I Henriksen, C Engelhard, AM Frøbert, M Schrölkamp, A Friebe, MT Overgaard, SS Poulsen, JF Rehfeld, J Nielsen, O Fröbert, C Scheele, A Feizi, S Nielsen

## Abstract

The brown bear (*Ursus arctos*) hibernates to survive cold winters without access to food. It builds enormous subcutaneous fat stores during summer and relies on it for energy during winter. Remarkably, the weight loss during winter occurs without muscle loss despite inactivity. Studying brown bear biology can therefore provide insights for improving human health in obesity and weight loss treatments. We here investigate subcutaneous adipose tissue biopsies obtained during summer and winter from free-living brown bears. During winter, a signature of genes involved in food intake and digestion is upregulated. Among these are several regulators of satiety, substrate transport and lipid metabolism. Interestingly, in humans these genes are enriched in distinct metabolic organs including, brain, intestine, stomach, liver and even salivary glands. We focused on the satiety brain/intestinal hormone cholecystokinin (CCK), which we demonstrate is produced in adipocytes, accompanied by an upregulation of the CCK receptor CCKBR. Importantly, CCK was undetectable in the circulation during winter and presence of sensory neurons suggest a neuronal feedback mechanism within the adipose tissue. Using RNA sequencing, we predict additionally 537 secreted proteins to be seasonally regulated, 37 of which could be confirmed with plasma proteomics. In conclusion, we propose that brown bears have developed a strategy of healthy fat burning and satiety regulation through an adipose tissue-contained mechanism which includes digestion factors and satiety mediators to provide safe energy turnover during hibernation-dependent weight loss.

## Introduction

Physical inactivity and excess of caloric intake in humans are major contributors to the global epidemic of obesity-related diseases including type 2 diabetes. Where excess weight in humans is associated with a wide range of comorbidities, the brown bear (*Ursus Arctos*) has adapted to seasonal fluctuations in food availability by undergoing yearly cycles of obesity and inactivity. As such, brown bears, shifting between an awake state and a periodic sleeping state have received attention as a fascinating model for comparative mammalian physiology and a promising means to study healthy obesity^1–8^. Characteristically, brown bears will hibernate for up to 7 months of the year^9^, while relying on stored adipose tissue as their sole source of energy and water^10^. Hibernation in bears is preceded by a state of hyperphagia during which the bears will gain of up to 30% of their spring body weight^11^.In Sweden this weight gain is predominantly based on consumption of berries^12,13^. In contrast with the anabolic hyperphagic phase, brown bears enter a prolonged state of hypometabolic fasting and physical inactivity when winter begins and they enter their sleeping phase. During hibernation, bears reduce their body temperature from 37°C to ∼33°C, decrease their metabolic rate to ∼25% of normal rate, slow their heart and respiratory rates, and enter a state of reversible insulin resistance^14^. Despite their extreme increase in adiposity and their low levels of physical activity bears are also seemingly protected from muscle atrophy, osteoporosis and cardiovascular diseases during hibernation^14^, whereas humans undergoing a similar scenario would be expected to show greatly increased risk of metabolic perturbation and atrophy of lean tissues^15^. The underlying mechanisms behind the protective properties of hibernating bears have been addressed in several studies^3,15^; in particular, adipose tissue of brown bears has received attention as a potential source of protective factors, due to its rapid changes in morphology^16^ and its’ role as the primary energy source during hibernation.

Adipose tissue has been established as an endocrine organ, secreting factors that regulate whole body metabolism^17^. In line with this, a plasma proteomics study revealed major season-dependent differences in brown bear blood^18^. However, it is not known if any of these factors are secreted from adipose tissue or whether novel factors not yet identified can be secreted from adipose tissue. To investigate how bear adipose tissue can contribute to a metabolically healthy profile during hibernation, we used a multi-omics approach to identify potential secreted factors from adipose tissue of free-range hibernating bears in both summer and winter and compared with expression levels in cultured bear adipocytes.

We identify 537 genes predicted to be secreted from bear adipocytes including high levels of leptin during winter. Surprisingly, we also find the gut satiety hormonecholecystokinin (CCK) is expressed and locally retained in the adipose tissue during winter. We therefore propose that brown bears have adapted to extended fasting by secreting CCK from their adipose tissue, providing an alternative source of satiety regulation, accompanied by elevated levels of leptin. Moreover, we characterize the seasonal metabolic properties of the bear adipose tissue, focusing on browning capacity and secretory pathways, and explore how these findings relate to humans.

## Results

### Bear winter adipose tissue is morphologically and transcriptionally distinct from summer adipose tissue

Blood and inguinal adipose tissue samples were collected from 3 male and 2 female free-ranging, 2-year-old subadult anesthetized brown bears during hibernation (February, 2014). Paired samples from the same animals were collected in the animals’ active state (July, 2014) (**Fig. 1A**). The body weight of the bears at the time of sampling are shown in **Fig. 1B**. The morphology of the adipocytes varied greatly between seasons; notably, adipocyte size was much larger in winter biopsies, whereas summer biopsies were characterized by smaller adipocyte size and more pronounced extracellular matrix formation (**Fig. 1C**).

**Figure 1.**
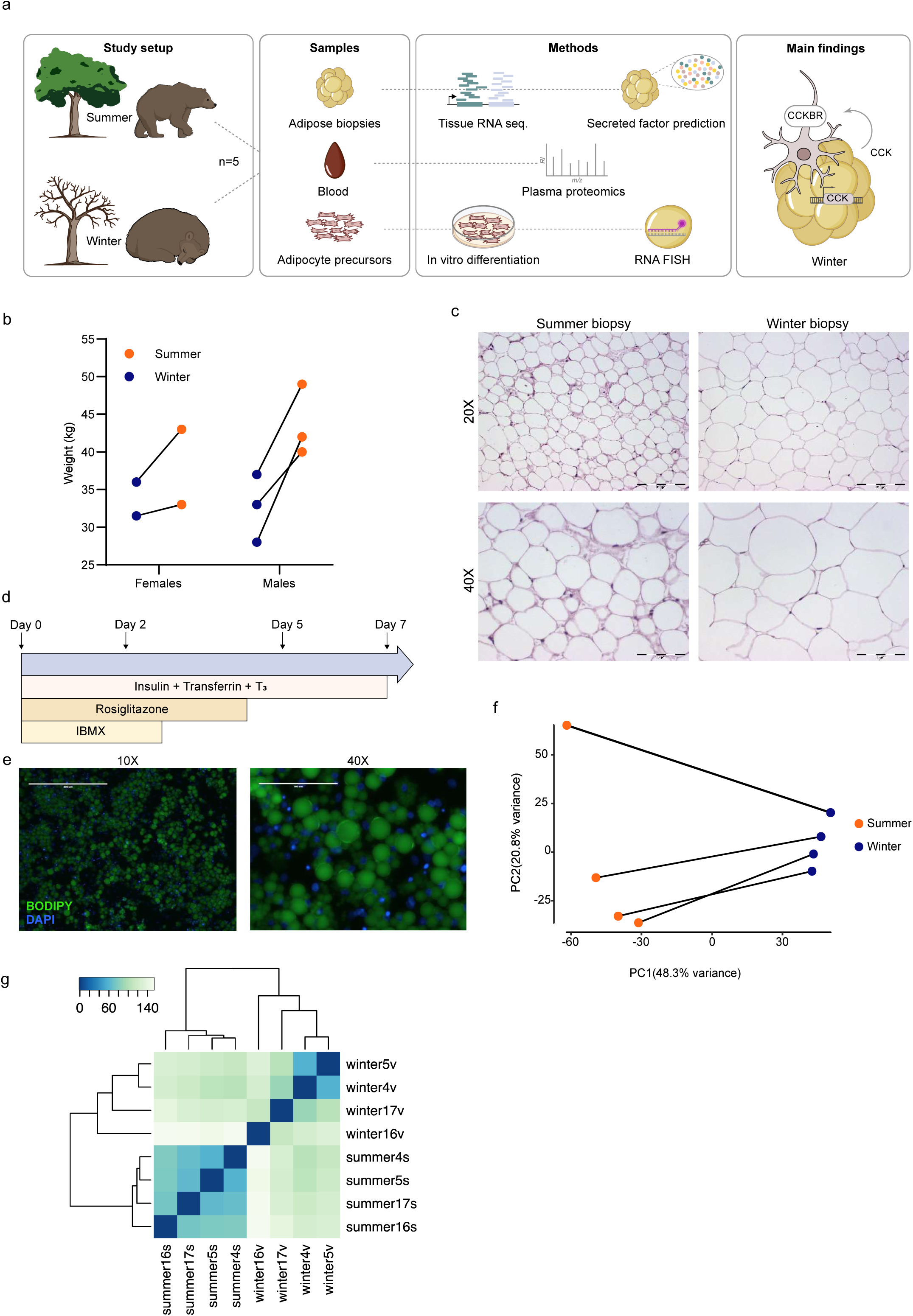
Morphology of tissue and cultured bear adipocytes. **a**. Overview of the study outline, methods and main results in this manuscript. **b**. Sex and weight of animals included in the study. **c**. Hematoxylin and eosin staining of adipose tissue sections taken during summer or winter at 20X (scale bar = 200 μM). or 40X (scale bar = 100 μM). **d**. Timeline and compounds used for in vitro differentiation of isolated bear preadipocytes. **e**. Representative image of in vitro cultured bear adipocytes at day 7 of differentiation; cells were fixed in 4% paraformaldehyde and lipid droplets were stained with BODIPY 505/515 (green) and nuclei were stained with DAPI (blue). **f**. Principal component analysis and correlation matrix (g) based on all detected genes in RNA sequencing dataset describing the differences between brown bear adipose tissue in the winter (blue) and summer (orange). **g**. A heatmap of sample-to-sample distances.

While this appears in contrast with the fact that the weight of the bears was lower in winter compared with summer (**Fig. 1B**) it may be explained by the young age of the bears in this study. From birth to until age 3 brown bears double in size each spring and continue to grow until 8-10 years of age^11,19,20^. The higher weight observed in summer, despite smaller adipocytes, may therefore be attributed to somatic growth in bears rather than solely to the expansion of adipose tissue.

Preadipocytes isolated from the biopsies were cultured in vitro following an abbreviated version of a human adipocyte differentiation protocol^21^ (**Fig. 1D**), and readily differentiated into multilocular, lipid-filled adipocytes (**Fig. 1E**).

To compare the transcriptional profiles of bear adipose tissue between summer and winter we performed Illumina RNA-sequencing on the adipose tissue samples (1 sample was excluded due to degraded RNA based on low RIN score). Principal component analysis (PCA) of the transcriptomes showed distinct clustering according to season (**Fig. 1F**), demonstrating that seasonal effects have global impact on bear adipose gene expression. A heatmap of sample-to-sample distances was used as an additional quality control (**Fig. 1G**)

### Food intake-related pathways are enriched in adipose tissue of hibernating bears

We measured global gene expression in winter adipose tissue compared with summer and found 3768 genes to be differentially expressed (DE) in winter samples. Surprisingly, the gene that showed the highest upregulation in winter biopsies was *CCK* (**Fig. 2A&B**), a peptide hormone primarily described as a gut hormone and neurotransmitter with effects on feeding and digestion^22^. Accordingly, gene set enrichment analysis (GSEA) using human gene ontology (GO) terms as a reference showed enrichment of several feeding-related processes driven by altered gene expression in winter biopsies (**Fig. 2C**), whereas processes related to connective tissues including cartilage development and bone metabolism were enriched in summer biopsies (**Fig. 2C**).

**Figure 2.**
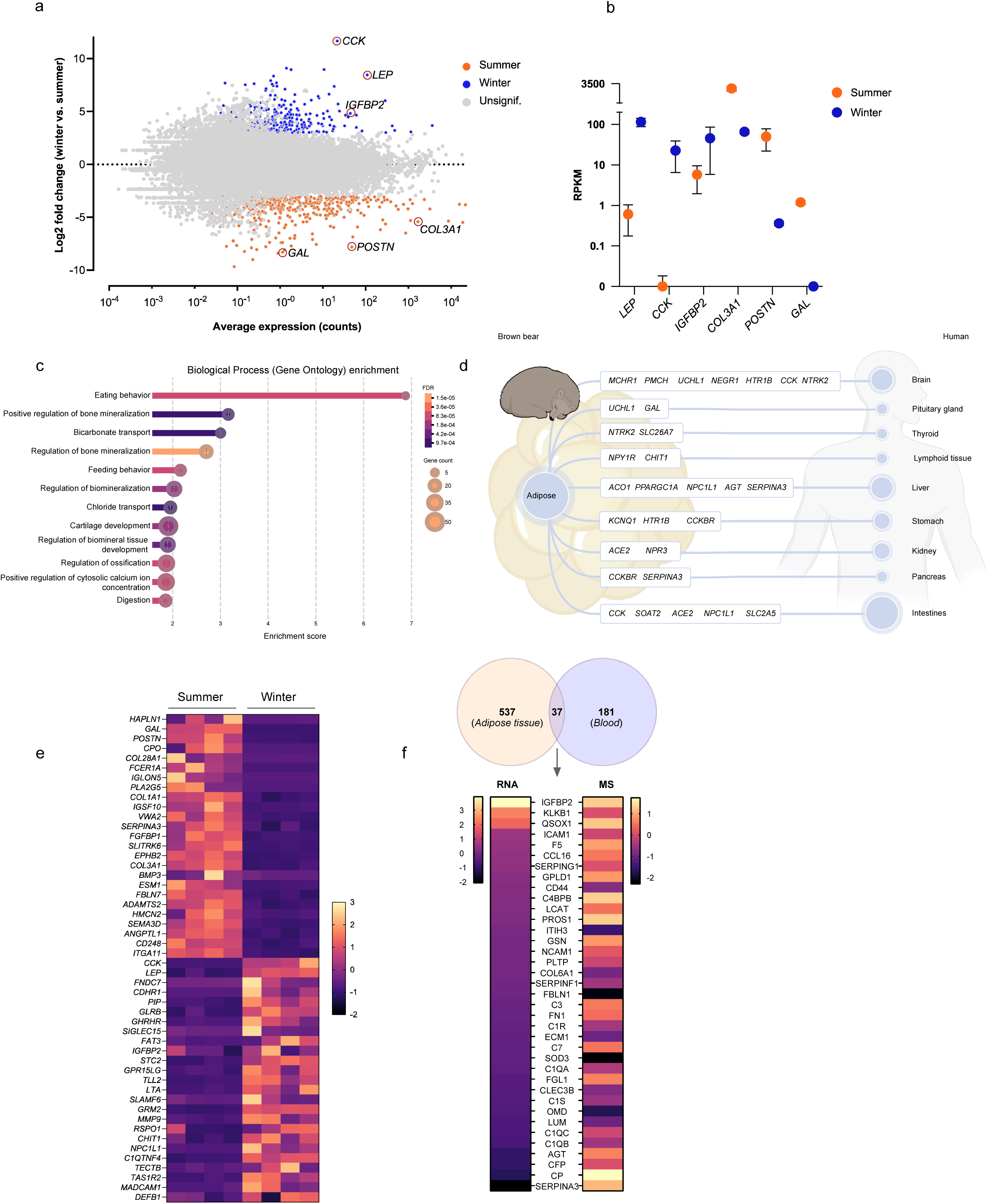
Transcriptomes of winter and summer brown bear adipose tissue. **a**. MA plot of differentially expressed genes between winter and summer; positive log2 fold changes are upregulated in winter and negative log2 fold changes are upregulated in summer (FDR < 0.01). **b**. CPM normalized gene expression of the three upregulated genes in winter (*LEP*, *CCK* and *IGFBP2*) or summer (*COL3A1*, *POSTN* and *GAL*) (FDR < 0.01). **c**. Gene set enrichment analysis of DE genes upregulated in winter versus summer. **d**. Schematic illustration of genes driving winter enrichment of feeding-related pathways in **c**. **e**. Heatmap visualizing seasonal differences in predicted secreted factors from bear adipose tissue (top 25 most upregulated genes in either season out of 537). **f**. Schematic illustration of overlap between DE genes detected by RNA sequencing and plasma proteomics.

During hibernation, brown bears restrict from food intake for months and the digestive tract is thus inactive and energy is provided solely from the adipose tissue. Interestingly, we observed a striking regulation of genes involved in digestion, eating and feeding behaviour in the brown bear adipose tissue during hibernation. In humans, these genes are expressed in multiple tissues involved in energy metabolism and regulation of food intake including pancreas, stomach, intestines, gut, liver, kidney and brain (**Fig. 2D**). The function of these factors included substrate transport (*SLC26A7*, *SLC2A5*) and intestinal absorption (*NPC1L1*, *SOAT2*) but most strikingly were the expression of several factors described to mediate satiety and food intake in humans. *LEP* was highly upregulated, consistent with its known role in humans where leptin regulates food intake through an adipose to brain-signaling axis. More surprisingly however, the satiety-mediating gut-hormone *CCK* and its receptor *CCKBR* were also upregulated in the bear adipose tissue while the orexigenic peptides galanin (*GAL*) and Pro-Melanin Concentrating Hormone (*PMCH*) and its corresponding receptor (*MCHR1*) were strongly downregulated (Supplementary table S1). In line with this, other obesity-related genes including *NEGR1*, *HTR1B*, *NPYR1* and *NTRK2* also showed differential expression in winter biopsies (Supplementary table S1), indicating a concerted regulation of satiety signaling in bear adipose tissue during the fasted winter period.

In line with previous reports^23,24^ *LEP* and *IGFBP2* expression were also upregulated in winter adipose tissue, while summer biopsies were characterized by upregulation of connective tissue and extracellular matrix (ECM)-related genes including *COL3A1* and *POSTN* (**Fig. 2A & 2B**).

To identify secreted factors in bear winter adipose tissue, we filtered all DE genes through SignalP 5.0 to detect genes harboring a secretory signal peptide^25^ thereby predicting these genes to be secreted. Clustering analysis of secreted DE genes showed a distinctly differential pattern (**Fig. 2E**), and GSEA showed overrepresentation of pathways related to ECM organization, ossification and blood vessel formation in summer biopsies and cell adhesion and migration winter biopsies (**Fig. S2A&B**).

To confirm the presence of secreted DE factors in the bears’ circulation we re-analyzed a mass spec (MS)-based proteomic analysis of bear plasma previously published from our group^26^. Interestingly, 37 out of 537 DE factors predicted to be secreted identified through adipose tissue RNA-sequencing were also detected by the MS-based approach on bear plasma (**Fig. 2F**), and although some of the overlapping genes showed opposing changes including Angiotensinogen (AGT), serpin family A member 3 (SERPINA3) and Ceruloplasmin (CP); **Fig. 2F**) these data suggest that the bear adipose tissue could be the source for the secreted factors. In contrast with being among the most upregulated genes detected by RNA-sequencing, leptin and CCK could not be detected by the used MS (**Fig. 2F**). This is most likely due to smaller size proteins are filtered away from the protein lysate, which is described in previous study^26^.

### Cholecystekinin is expressed in bear adipose tissue during winter but not summer

While the CCK receptors have been found to be present in adipose tissue in rats^27^, expression of CCK has not been previously reported in adipose tissue or adipocytes. To assess if bear CCK represents a functionally distinct peptide we compared the amino acid sequences of human and bear preproCCK, which were highly similar (**Fig. 3A**). Importantly, the discrepant amino acids were all located distally to the C-terminally active site of CCK (CCK-8)^22^, which was fully conserved between species, and are therefore unlikely to represent a functionally relevant difference. Moreover, comparison of the bear CCK locus in summer and winter adipose biopsies showed that a single 3-exon transcript was expressed in winter only, suggesting that differential transcript usage could not explain the season-dependent change in *CCK* expression levels (**Fig. 3B**).

**Figure 3.**
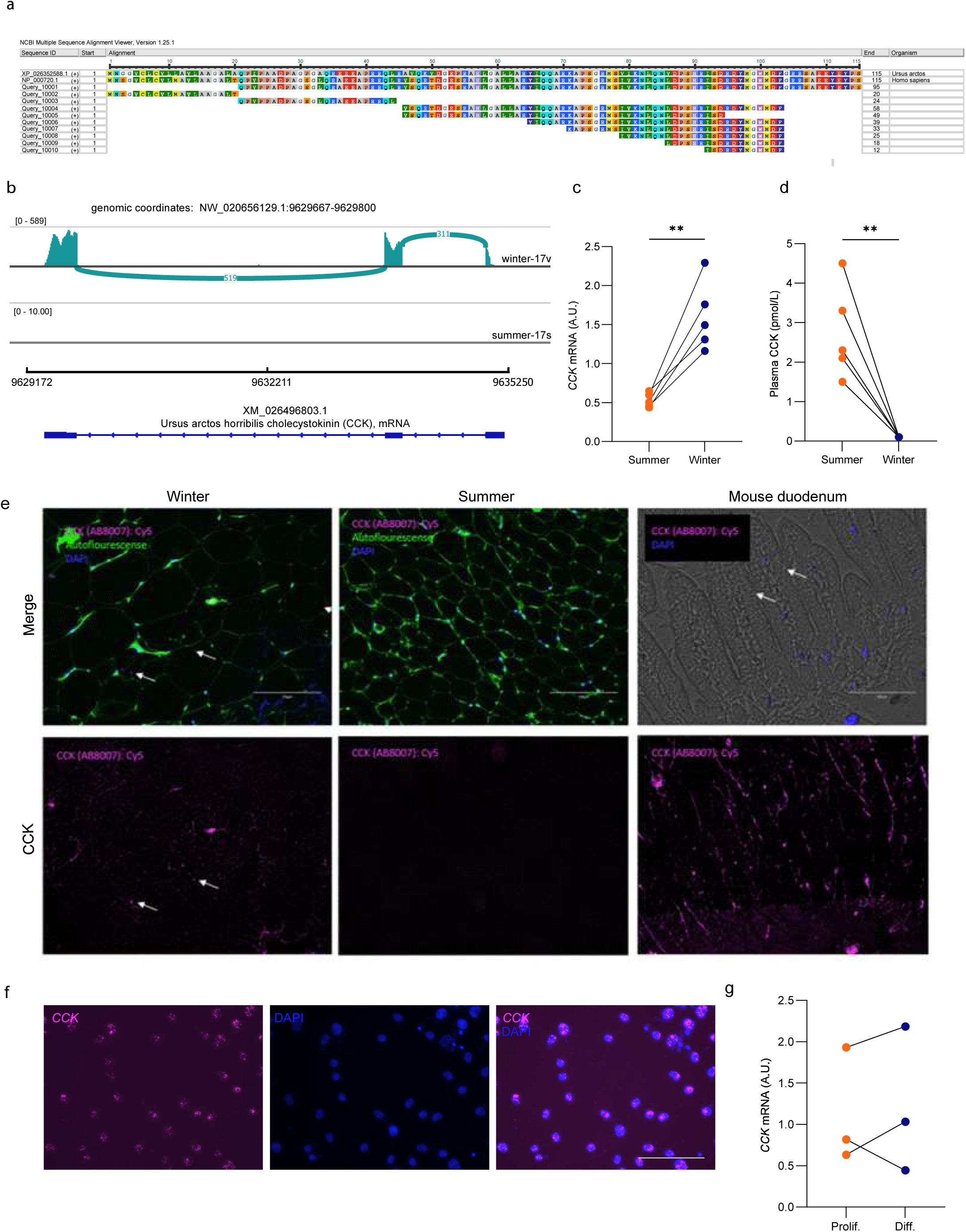
Expression of cholecystokinin in brown bears in winter versus summer. **a**. Alignment of bear CCK protein sequence (top) versus human CCK isoforms. **b**. Sashimi plot showing bear CCK exon usage in winter versus summer. **c**. qPCR quantification of CCK mRNA expression in bear winter versus summer adipose tissue. **d**. Plasma protein levels of CCK in winter versus summer. **e**. Immunofluorescent labelling of CCK protein (magenta) in paraffin-embedded bear adipose tissue sections from summer or winter; tissue autofluorescence in the GFP spectrum indicates cell morphology (green); nuclei were stained with DAPI (blue). Mouse duodenum was included as a positive control (scale bar = 150 μM). **f**. Fluorescent in situ hybridization of CCK mRNA (magenta) in cultured bear adipocytes at day 7 of differentiation. Scale bar indicates 75μm.

Accordingly, the observed increase in adipose *CCK* expression in winter compared with summer biopsies was confirmed by qPCR (**Fig. 3C**). To confirm the presence of the CCK peptide in winter adipose tissue, we performed immunofluorescent staining of adipose tissue sections from summer and winter biopsies. In line with the seasonal *CCK* mRNA expression patterns, the CCK peptide was only detectable in winter sections (**Fig. 3D**), while plasma CCK levels were markedly decreased in winter compared with summer (**Fig. 3E**). This suggests that CCK is likely released from other tissues such as the gut in response to feeding when the bears are active, whereas adipose-derived CCK may have local rather than systemic effects. To further confirm that adipocytes were indeed expressing the CCK transcript we performed RNA fluorescent in situ hybridization (FISH) analysis in *in vitro* cultured fully differentiated bear adipocytes. Intriguingly, most of the cells were *CCK*–positive with the transcript being exclusively located to the nucleus (**Fig 3F**). In vivo, duodenal cells secrete CCK in response to byproducts of feeding, such as amino acids and lipids; to identify a potential signal leading to translocation of the *CCK* transcript from the nucleus to the cytosol for translation we therefore exposed the cells to factors that could mimic a post-prandial situation in the adipocyte cell cultures. The adipocytes were consequently treated with varying concentrations of amino acids including tryptophan, phenylalanine and glutamine, as well as different lipolytic factors including norepinephrine, IL-6, atrial natriuretic peptide and leptin; however, we were not able to detect any changes in subcellular *CCK* location (data not shown). Furthermore, qPCR analysis of cultured bear adipocytes showed no differences in CCK expression during preadipocyte proliferation compared with fully differentiated cells (**Fig. 3G**).

Collectively, these data show that bear adipose tissue expresses CCK exclusively during winter, while other tissues secrete it during summer. Additionally, bear adipose tissue may serve as a reservoir for CCK mRNA during hibernation.

### Adipose tissue of hibernating bears shows increased expression of afferent markers but reduced browning

We hypothesized that CCK may only be secreted locally in bear winter adipose tissue to act in a paracrine manner. In the gut, CCK can signal through vagal afferent neurons expressing the CCK-A receptor to mediate feeding suppression^22^. We therefore measured the expression of the CCK receptors in adipose tissue biopsies from summer and winter. Interestingly, while the CCK-A receptor was not detected in either season, the CCK-B receptor was expressed in both seasons and upregulated in winter (**Fig. 4A**). In line with this, we detected the protein expression of the sensory neuronal marker calcitonin gene–related peptide (CGRP), further supporting the notion that local CCK signaling may take place in bear adipose tissue (**Fig. 4B**).

**Figure 4.**
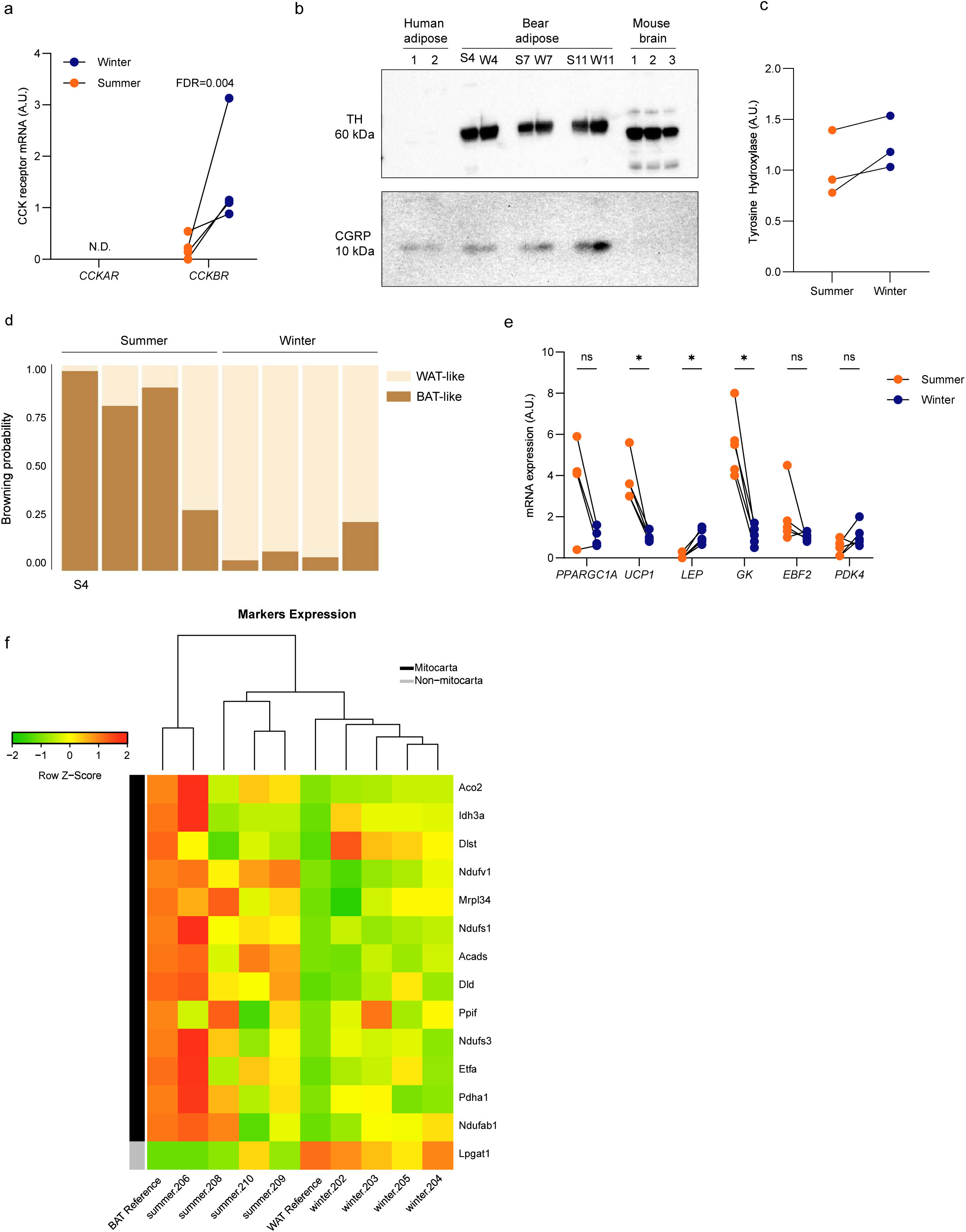
Expression of bear adipose tissue neuronal markers and brown fat genes. **a**. qPCR quantification of CCKAR and CCKBR mRNA expression in bear winter versus summer adipose tissue. **b**. Immunoblotting of Tyrosine hydroxylase (TH), and Calcitonin gene-related peptide (CGRP) in human subcutaneous adipose tissue, bear adipose tissue from winter or summer, and in mouse brain. **c**. quantification of bear TH protein expression. **d**. ProFAT analysis of bear winter or summer adipose tissue transcriptomes. **e**. qPCR quantification of brown fat-related and adipogenic gene expression in bear winter versus summer adipose tissue. **f**. Heatmap of seasonal differences in mitochondria-associated genes in bear adipose tissue compared with a brown or white adipose reference gene set. P<0.05

Moreover, tyrosine hydroxylase (TH) was also expressed in both summer and winter biopsies (**Fig. 4B**), suggesting the presence of sympathetic neurons in bear adipose tissue. CCK signaling has been implied in the regulation of brown adipose activity^28,29^, we therefore examined the extent of brown fat gene expression in bear adipose biopsies. Interestingly, ProFat analysis^30^ showed a substantially higher browning probability in summer adipose samples compared to winter (**Fig 4D**), and qPCR analysis of central brown fat genes including uncoupling protein 1 (UCP1) showed a decrease in winter adipose biopsies (**Fig. 4E**). In line with this, summer adipose biopsies showed a more brown adipose tissue (BAT)-like profile when comparing with human Mitocarta genes^31^, suggesting that bear adipose tissue adopts a more brown-like gene expression profile during the summer season (**Fig. 4F**). However, it should be noted that the ProFat and Mitocarta genes are dominated by mitochondrial genes, and that the algoritms have not been trained on adipose tissue from bears. A high expression of mitochondrial genes would also be required for the extreme adipogenesis occurring in the bears during summer and might thus be less related to browning.

Collectively, these data suggest that bear adipose tissue is both efferently and afferently innervated, supporting the notion that CCK may act in a paracrine manner, and moreover, that bear BAT activity may be higher in summer than in winter.

## Discussion

We here show that adipose tissue from hibernating brown bears activates a distinct transcriptional profile comprising a range of neuronal and satiety-regulating factors, including CCK and the CCKB receptor, suggesting the presence of a fat-to-brain satiety axis within the adipose tissue during hibernation. Moreover, our sequencing data shows that in the absence of feeding brown bear adipose tissue turns on expression of genes which in humans associate with digestive organs including brain, intestines, liver, stomach, and kidney, indicating that a shift towards a centralized function of these organs takes place in the bears’ adipose tissue during hibernation. Since adipose tissue serves as the primary energy source during the prolonged state of fasting during winter, this transcriptional mirroring of digestive organs may reflect that adipose remains an active metabolic hub while the canonical digestive organs reduce their activity. This functional shift underscores the highly adaptive capacity of adipose tissue, allowing it to compensate for the reduced metabolic activity of other organs, while ensuring the bear’s energy demands and appetite control are maintained throughout hibernation.

Previous studies have compared seasonal gene expression across different metabolic tissues in brown bears, with adipose tissue consistently showing the highest percentage of differentially expressed (DE) genes during hibernation. For instance, Jansen et al. reported approximately 6,000 DE genes in adipose tissue^3^, while we identified 3,768 DE genes in our study. Notably, our research focuses on free-ranging brown bears, in contrast to the captive bears studied previously. The higher number of DE genes reported by Jansen et al. may reflect differences in data processing methods and the distinct biological contexts of the two models.

Furthermore, the lack of overlap in winter adipose gene set enrichment analysis (GSEA) terms between the two studies may also be influenced by the differences between captive and free-ranging life.

In humans, food intake and satiety are regulated by a complex network of hormones spanning multiple tissues, and euroendocrine signaling from the alimentary tract to the brain in response to feeding is central to this process^32,33^. Our data suggest that, during hibernation, brown bear adipose tissue integrates these signals through an adipose-neuronal mechanism, compensating for the absence of gut-derived fed-state signals.

Moreover, our findings suggest that adipose tissue from hibernating brown bears may hold promising candidates for the development of novel polypharmacologic drugs designed to target multiple pathways involved in food intake regulation. Current incretin-based weight loss drugs, while effective in reducing body weight in humans with obesity, often lead to excessive lean mass loss^34^, which can have long-term health consequences. In contrast, hibernating brown bears maintain lean mass during prolonged fasting and inactivity, and it is possible that this ability stems from their adipose tissue’s integration of energy expenditure with energy consumption and satiety regulation, enabling fine-tuning of weight management without hunger or loss of lean mass. Beyond feeding behavior regulation, several of the satiety-related factors showing altered expression in winter adipose tissue—including galanin, leptin, CCK, PMCH, and NPY1R—are also implicated in sleep regulation^35–38^. This raises the intriguing possibility that bear adipose tissue contributes not only to hunger suppression but also to sleep regulation during hibernation.

Our findings further position the brown bear as a valuable and novel animal model for studying human metabolism. Mice are widely used in metabolic research, but their biology—and particularly their metabolic processes—differs significantly from humans limiting their translational relevance. In contrast, the remarkable seasonal adaptations observed in free-ranging brown bears reflect naturally evolved mechanisms that enable survival in their environment. While the lack of option to do interventions when studying bears is a limitation, these evolutionary adaptations nevertheless offer valuable insights into metabolic regulation that may be harnessed to develop treatments for metabolic diseases that current models fail to address effectively.

Understanding these evolutionary mechanisms provides a foundation for novel therapeutic strategies, paving the way for more effective interventions in human metabolic health.

## Conclusions

- Brown bears have adapted to switching on CCK expression in their adipose tissue stores during hibernation,
- Although CCK is not secreted from bear adipose systemically, the presence of afferent neurons and expression of the CCKR suggest that CCK may act as a locally secreted neuroendocrine factor
- Bear adipose tissue may therefore act as an alternative source of satiety signals during long term fasting

## Materials and Methods

### Capturing of bears

Blood and subcutaneous adipose tissue were collected from immobilized free□ranging brown bears (fitted with GPS□collars) during hibernation in winter (early February) and, from the same bears, during the active period in early summer (late June/early July) in Dalarna, Sweden. The adipose tissue was taken from the inguinal fat depot during general anesthesia. All captures were approved by the Swedish Ethical Committee on Animal Research (C212/9 and C47/9) and the Swedish Environmental Protection Agency and follow the general principles of Public Health Service Policy on Humane Care and Use of Laboratory Animals applicable to wildlife. Sampling was done by the Scandinavian Brown Bear Research Project (https://www.brownbearproject.com) according to established protocols (Arnemo et al. 2012)

### Adipose Tissue sectioning and staining

Bear adipose tissue biopsies were fixed in 4% formalin buffer (Cellpath, Newtown, UK) and embedded in paraffin. 3 μm sections were heated at 60°C for 60 min, deparaffinized in Tissue□Clear (Sakura Finetek Europe, Alphan, the Netherlands) and rehydrated. CCK staining’s were performed using an antibody AB8007 developed by Jens Rehfeld, described here (PMID). The antibody was diluted 1:6000 in HBSS and incubated with the bear tissue slides overnight at 5°C. The slides were washed twice with DPBS and the incubated with goat anti rabbit Cy5 (A10523) for 30 min at room temperature. To visualize the nuclei, a co-staining with Nucblue ReadyProbes (R37605, Invitrogen; 2 drops pr ml) was performed for 20 min at room temperature. The fluorescence signals were detected with DAPI and Cy5 light cubes using a EVOS M5000 microscope. Light intensity was adjusted to the duodenum slide (positive control) so we achieved minimum background. The same light intensity settings were used for both the winter and summer slides.

### RNA FISH

In vitro differentiated bear adipocytes derived from the inguinal adipose depot were fixed on days 4 and 6 of differentiation with 10% neutral buffered formalin (HT501128-4L, Sigma) for 30□min, dehydrated and stored in 100% ethanol until the staining procedure. In situ hybridization was performed using RNAscope Multiplex Fluorescent Detection Kit v2 and RNAscope 4-Plex Ancillary Kit for Multiplex Fluorescent Kit v2 (323110 and 233120, ACDbio). RNAscope manual assay probes were designed and produced by Advanced Cell Diagnostics. Nuclei were stained with NucBlue. RNA targets were visualized using an EVOS imaging system. The RNA targets were hybridized with bear specific CCK RNAscope probes (Cat No. 895641) and then labelled with Opal 690 (FP1497001KT; Akoya Biosciences). The fluorescence signals were detected with DAPI and Cy5 light cubes.

### Culturing of cells

Due to an extraordinary ability to accumulate lipids, a modified protocol for culturing primary cell cultures was needed. A previously described protocol was used as template to establish the modified protocol^39^. After obtaining the biopsy from the inguinal fat depot in the field, the biopsy was placed in a 50 ml tube containing DMEM/F12 (Gibco) with 1% penicillin/streptomycin (PS; 15140122, Life Technologies; 10,000 U/mL).

Tubes were kept on ice during the transport from the field to the cell-lab. Adipogenic progenitor cells were isolated from the stromal vascular fraction of the biopsies on the day they were obtained. Biopsies were digested in a buffer containing 10 mg collagenase II (C6885-1G, Sigma) and 100 mg BSA (A8806-5G, Sigma) in 10 ml DMEM/F12 for 20–30 min at 37 °C while gently shaken. Following digestion, the suspension was filtered through a cell strainer (70 μm size) and cells were left to settle for 5 min before the layer below the floating, mature adipocytes was filtered through a thin filter (30 micron). The cell suspension was centrifuged for 7 min at 800 g, and the cell pellet was washed with DMEM/F12 and then centrifuged again before being resuspended in DMEM/F12, 1% PS, 10% fetal bovine serum (FBS) (Life technologies) and seeded in a 25 cm2 culture flask. Media was changed the day following isolation and then every second day until cells were 80% confluent; at this point, cultures were split into a 10 cm dish (passage 0).

Progenitor cells were expanded in proliferation media consisting of DMEM/F12, 10% FBS, 1% PS and 1 nM Fibroblast growth factor-acidic (FGF-1) (ImmunoTools). Cells were grown at 37 °C in an atmosphere of 5% CO2 and the medium was changed every second day. Adipocyte differentiation was induced two days after preadipocyte cultures were 100% confluent by addition of a differentiation cocktail consisting of DMEM/F12 containing 1% PS, 0.1 μM dexamethasone (Sigma–Aldrich), 100 nM insulin (Actrapid, Novo Nordisk or Humulin, Eli Lilly), 200 nM rosiglitazone (Sigma–Aldrich), 540 μM isobutylmethylxanthine (Sigma–Aldrich), 2 nM T3 (Sigma–Aldrich) and 10 μg/ml transferrin (Sigma–Aldrich). After three days of differentiation, isobutylmethylxanthine was removed from the cell culture media, and after additional an day rosiglitazone was removed from the media for the remaining 3 days of differentiation.

### qPCR

Total RNA isolation from adipose tissue biopsies or cultured adipocytes was performed using TRizol reagent according to the manufacturer’s protocol. RNA was dissolved in nuclease-free water and quantified using a Nanodrop ND 1000 (Saveen Biotech). Total RNA (0.25 μg) was reverse-transcribed using the High Capacity cDNA Reverse Transcription Kit (Applied Biosystems). cDNA samples were loaded in triplicate and qPCR was performed using Real Time quantitative PCR, using the ViiA7 platform (Applied Biosystems). Relative quantification was conducted using SYBRgreen fluorescent dye (A25742, Applied Biosystems). All procedures were performed according to the manufacturer’s protocol. Target mRNA expression was calculated based on the standard curve method. Primers targeting CCK mRNA was designed using ProbeFinder version 2.50 (http://www.universalprobelibrary.com, Roche Applied Science).

### Primers

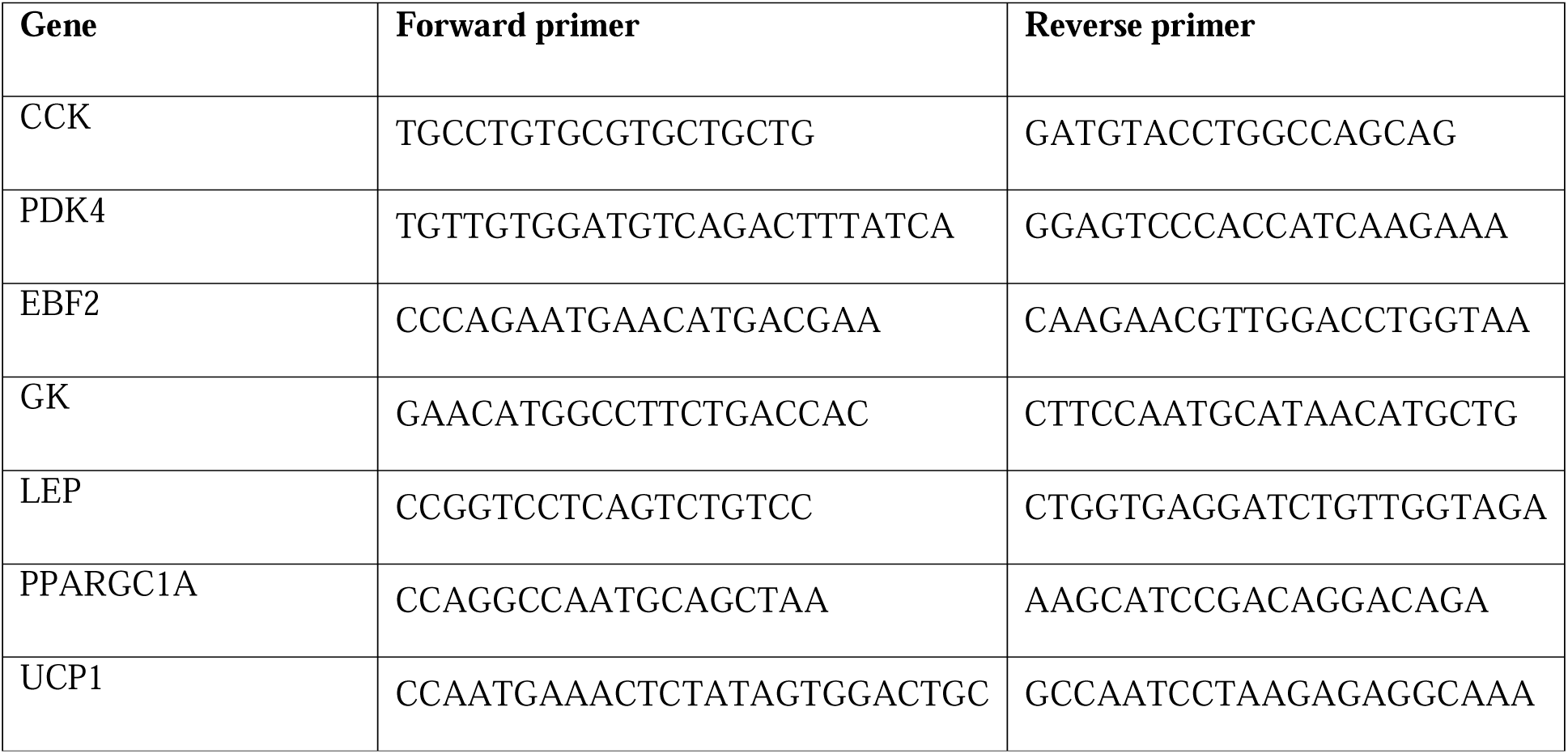

#### RNA-sequencing

The pair-end fastq files (raw reads) from the 10 samples were pre-processed by Trimmomatic^40^ to remove adaptors, leading low quality or N bases (below quality 3), quality trimming (4-base wide sliding window and quality cut-off of 15), and removing reads below 36 bases long. The remaining reads were mapped and quantified by STAR aligner against the brown bear genome as a reference^41^. MultiQC was used for quality control of the reads and alignment. We used DESeq2 R packages^42^ for DE analysis and MA plot generation. Gene counts were normalised using DESeq2’s regularized logarithm transformation and principal component analysis was performed after centering over the different samples. We used IGV viewer for visualizing the read counts^43^. DE genes were screened for signal peptides using Signal.P 5.0^25^. To generate a heatmap of predicted secreted genes, gene counts were first normalized using DESeq2’s regularized logarithm transformation followed by centering and scaling over the different samples by the root-mean-square (Z score). Subsequently, genes were clustered by hierarchical clustering according to their euclidean distance using the complete linkage method. For GSEA analysis, all DE genes (FDR P value< 0.05; 3768 genes in total) were uploaded to https://string-db.org/ along with their corresponding LOG2 fold change (no cutoff), using the ‘Proteins with Values/Ranks - Functional Enrichment Analysis’ function^44^. Human GO terms was used as a reference. Filtering resulted in 2777 genes queried. FDR stringency was set to ‘High’ (1 percent).

#### Protein expression

Brown bear inguinal adipose protein expression was assessed by western blotting. Primary antibodies were used at the following dilutions: CGRP (Cell Signaling D5R8F; 1:1000), tyrosine hydroxylase (Abcam ab112; 1:200). Primary antibodies were detected with either anti-rabbit or anti-mouse horseradish peroxidase-linked IgG (Dako) at a concentration of 1:5000 and imaged using Supersignal West Femto (Pierce). Data are expressed relative to total protein expression using stain free UV imaging (Biorad). Chemiluminescent signals were quantified using Image Lab version 5.2.1 software (Biorad).

#### CCK Measurements in blood

CCK was measured in brown bear EDTA-plasma with a RIA kit developed by Jens Rehfeld^45^.

### Proteomics

#### Analysis software and statistics for cell experiments

Data are represented as means□±□s.e.m. Statistical analysis was performed with GraphPad Prism version 10. A paired t-test was performed as indicated in the figure legends. N values are stated in the figure legends

## Acknowledgements

The laboratory procedures were performed at The Centre for Physical Activity Research which is supported by grants from TrygFonden (nos. 101390 and 20045). During the study period, the Centre of Inflammation and Metabolism was supported by a grant from the Danish National Research Foundation (no. DNRF55). S.N. was further supported by the Novo Nordisk Foundation (NNF20OC0061400). This work was also supported by the Novo Nordisk Foundation (NNF21SA0072102,) This study was supported by the Scandinavian Brown Bear Research Project, SBBRP (Norwegian Institute for Nature Research / https://bearproject.info), funded by the Norwegian Environment Agency and the Swedish Environmental Protection Agency. We thank the field capture team of the SBBRP and rangers from Region Gävleborg, Sweden for performing the bear captures and the veterinary team headed by Prof. Jon Arnemo responsible for anesthesia and sampling.

