## Supplementary figures for "Brown Bears activates a Satiety Hormone Cholecystokinin (CCK) pathway in adipose tissue during hibernation"

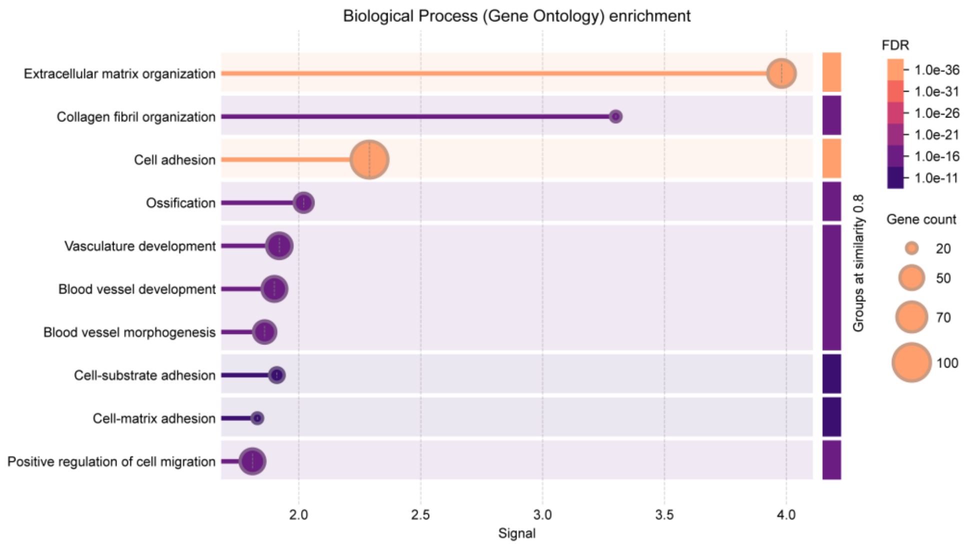


**Figure S2A:** Gene set enrichment analysis of summer-upregulated DE genes predicted to be secreted by SignalP.

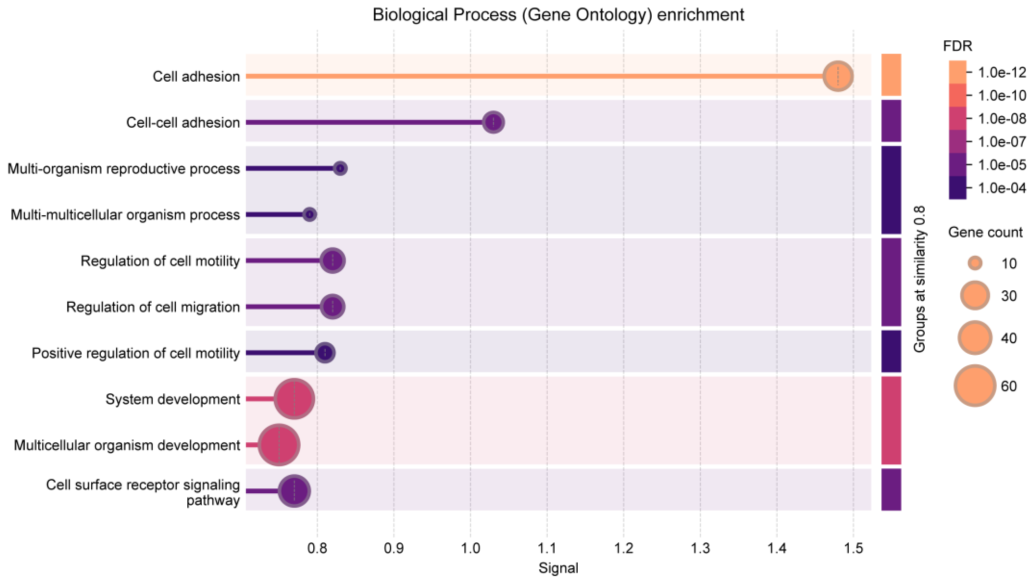


**Figure S2B:** Gene set enrichment analysis of winter-upregulated DE genes predicted to be secreted by SignalP.
